# A network model of path control

**DOI:** 10.64898/2026.10.02.756268

**Authors:** Mohammed Abdal Monium Osman, Wenyi Zhang, Cengiz Pehlevan, Rachel I. Wilson

**Affiliations:** Department of Neurobiology, Harvard Medical School; Kempner Institute for the Study of Natural and Artificial Intelligence, Harvard University; John A. Paulson School of Engineering and Applied Sciences, Harvard University; Center for Brain Science, Harvard University

## Abstract

We present a network model that enables a navigating agent to shape the spatial and temporal structure of its paths through its environment. The model is inspired by the insect fan-shaped body. In it, a recurrent neural network receives an input that encodes an agent’s velocity vector. Meanwhile, network activity determines the setpoint of a control loop that influences this velocity vector through motor outputs. The network thus guides future movement using memory of past movement. We illustrate how the network’s dynamics determine the trajectory features it extracts to produce the setpoint. We then relate the joint dynamics of network activity and the agent’s velocity vector to the structure of the resulting paths. This network can produce paths suitable for different forms of exploration. It can also stabilize movements initiated by lower-level systems. Our analyses illustrate how recurrent neural dynamics and memory can shape behavior when embedded in closed loops.

## 1 Introduction

The fan-shaped body (FB) of the insect central complex is a brain region downstream of the head direction system. It contains a set of neural arrays in which sinusoidal activity patterns encode vectors relevant to navigation [1–5]. Each array constitutes a “columnar cell type” of the FB (Fig. 1A). Activity across these arrays is coupled by highly structured, array-specific patterns of recurrent connectivity (Fig. 1B,C; [3]). Whereas feedforward networks of such arrays can implement arithmetic operations like vector addition and rotation (Fig. 1D,E; [1, 2]), it is not entirely clear what computations are performed by recurrent ensembles of sinusoidal arrays.

**Figure 1:**
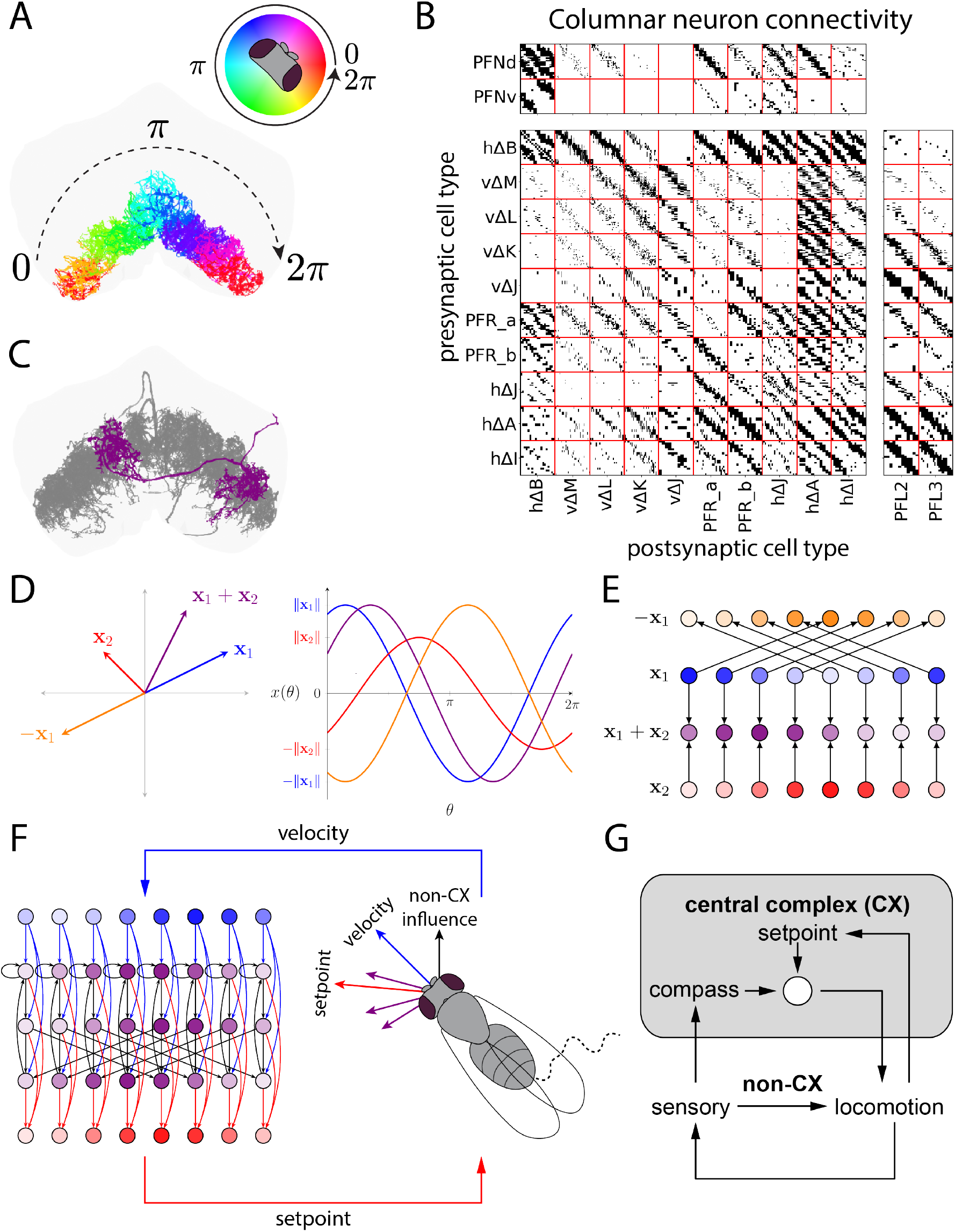
Columnar organization of the FB. (A) Neurons of a given columnar cell type tile the FB, subdividing it into columns that correspond to the 2*π* angular extent of compass directions. (B) Binarized connectivity of a subset of the columnar cells downstream of the network’s velocity input (PFNd/PFNv) and upstream of the network’s motor output (PFL2/PFL3). Note that within each block, neurons of a given pre- and post-synaptic type, sorted by their column, topographically connect with some characteristic strength. Each block consists of phase-preserving (diagonal) and/or phase-inverting (off-diagonal) connections. (C) Phase-inverting connections are permitted by the morphology of hΔ neurons, which contain arbors in pairs of opposing columns. (D) A two-dimensional vector can be associated with a sinusoidal activity pattern whose phase and amplitude match the vector’s direction and length, an encoding scheme called a phasor representation [2]. (E) Phasor representations can be instantiated by spatial activity patterns in arrays of columnar neurons, with simple feedforward motifs implementing operations like vector addition and inversion [1]. (F) The animal’s behavior and neural activity define a closed loop. In it, behavior influences neural activity through an input that encodes the animal’s velocity vector (blue), this input is processed by a recurrent network of sinusoidal arrays (purple), and network activity influences behavior through a setpoint signal (red) produced by summing the vectors encoded by these activity patterns. Finally, non-CX circuits independent of this loop influence the velocity vector in parallel (represented by the black arrow). (G) Block diagram schematizing the system’s control architecture.

To understand this network, we need to consider the inputs and outputs that couple network activity to behavior (Fig. 1F,G). One of the chief inputs to the FB is a sinusoidal array whose activity encodes the animal’s velocity vector, i.e. its current direction and speed of travel in the ground plane (PFNd/PFNv cells) [4, 5]. It has been proposed that the FB might use this input for path integration, the estimation of positional displacement from a time series of velocity measurements [6–10]. Path integration is commonly thought to underlie homing behavior in insects like bees and ants [11–13]. It has been suggested that a sinusoidal array could path-integrate a velocity input to produce a sinusoid that encodes position [14].

While path integration remains a plausible role for the FB, recent work in *Drosophila* suggests a more complicated picture. Based on recent connectomes [3], the FB actually contains many parallel pathways downstream of the velocity input, making the network more intricate than we might expect from minimal models of path integration (e.g. [14]). In addition, recent calcium imaging experiments have found no columnar cell types that encode a classical path integral; instead, many of the relevant cell types seem to encode lagged and smoothed versions of the velocity input [15–18], consistent with relatively transient, leaky integration dynamics. These results suggest that the FB may temporally filter its velocity input in more varied ways than classical path integration, but it remains unclear how these more transient forms of integration might facilitate navigation.

On the output side, FB activity contributes to locomotor control. Specifically, PFL2/PFL3 cells continuously compare the animal’s heading to a setpoint specified by activity in the FB; whenever the animal’s heading deviates from this setpoint, PFL2/PFL3 cells slow the animal’s forward speed and drive corrective turning toward the setpoint [15, 19]. This setpoint is a two-dimensional vector encoded by the summed activity patterns of multiple arrays (columnar cell types) in the FB. These arrays are themselves down-stream from the velocity vector input signal. Thus, the FB is part of a closed loop in which behavior (the animal’s velocity vector) influences neural activity, and vice versa.

Finally, it is also important to recognize that the FB does not completely specify locomotor behavior. Instead, behavior emerges from the combined influence of the FB and other circuits acting in parallel, such as lower-level sensorimotor systems that steer the animal toward or away from salient stimuli. It remains unclear how these parallel pathways are coordinated with the effects of the FB.

To explore the FB’s role in navigation, we developed a network model based on its architecture. In this model, a network of recurrently interacting sinusoidal arrays receives an external input that encodes an agent’s velocity vector. We first illustrate how the network’s recurrent connectivity determines the temporal filter that each array applies to the velocity input. This filtering allows the network to extract information about the agent’s trajectory over a variety of time scales and frequency bands, and it generalizes classical path integration to more varied forms of signal processing.

We then couple the agent’s locomotor behavior to network activity through a control policy based on that of the FB [6, 15, 19, 20], thus defining a closed neural-behavioral loop. Within our model, agent behavior is also influenced by a forcing function meant to capture lower-level sensorimotor pathways that are independent of the FB. Analyzing this neural-behavioral system, we relate the structure of the network and forcing function to the shapes of the agent’s paths through its environment. The resulting paths can combine meandering, oscillatory, and persistent movement, potentially supporting a variety of exploratory behaviors. The network can also stabilize movements initiated by the lower-level pathways, thereby enabling the agent to persistently move toward an intermittently sensed cue.

Based on this modeling, we propose that the FB may use its velocity input to extract and store the features of past movement needed to generate a path with a desired spatial structure. These results demonstrate how memory of past movement can be used to organize structured behavioral sequences. More generally, they illustrate how recurrent neural circuits embedded in closed loops can shape behavior.

## 2 Results

### 2.1 A network model inspired by the FB

The FB is divided into horizontal rows and vertical columns. Different rows (or “layers”) serve different functions [3]. In this study, we will focus on the intermediate layers of the FB, which contain the circuitry most directly implicated in spatial navigation [4–6, 15, 19]. Each layer is horizontally tiled by “columnar cells”. Each columnar cell type forms a complete set of tiles, and cells with adjacent horizontal positions are tuned to adjacent compass directions (Fig. 1A; [3]). We will model the FB as *N* populations of neurons, each representing a columnar cell type. Each neuron in this network model will be indexed by a cell type label *i* ∈ {1, …, *N*} indicating the population it belongs to and an angular position *θ* ∈ [0, 2*π*) indicating its column.

Within the FB, columnar neurons are coupled by recurrent connections whose strength depends on their respective cell types and relative positions (Fig. 1B; [3]). These connections come in two varieties: phase-aligned connections between neurons that share the same column, and phase-inverted connections between neurons that occupy opposing columns (columns separated by exactly half the horizontal extent of the FB) (Fig. 1B; [3]). These phase-inverting connections are enabled by a class of columnar cell types called hΔ (Fig. 1C; [3, 21, 22]).

Within our network model, the strengths of these connections are represented by two matrices: **W**^+^, 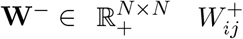 parameterizes the strength of the phase-aligned connections from neurons of cell type *j* to those of cell type *i*, and 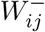 is defined analogously for phase-inverted connections. In our model, all these connections are non-negative, a choice motivated by experimental evidence that FB columnar neurons are excitatory [23, 24].

Next, we consider the input that informs this network about the agent’s behavior. In *Drosophila*, neurons in the intermediate layers of the FB receive input from cells (PFNd/PFNv cells; [4, 5]) that encode the animal’s velocity vector via a sinusoidal spatial pattern, an encoding scheme called a phasor representation (Fig. 1D,E; [1, 2]). We model this input as a spatial activity profile that encodes the current velocity **v**(*t*) ∈ ℝ^2^ of an agent moving through 2D space (Fig. 2A-C):

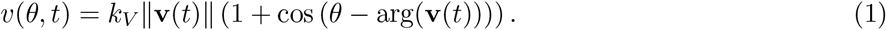

**Figure 2:**
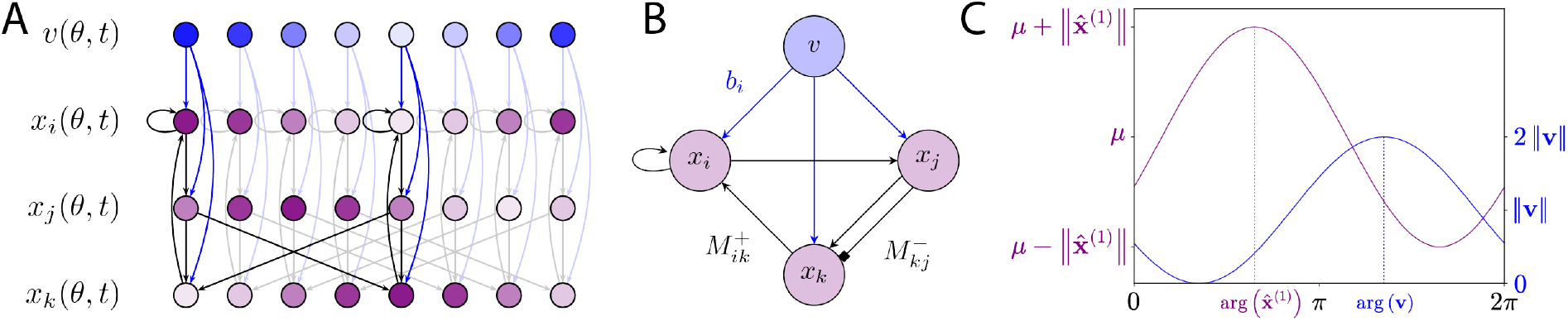
Overview of network model. (A) The network model consists of a set of neural arrays, each meant to represent a columnar cell type. Each array receives an external input that encodes the agent’s velocity. Neurons are recurrently coupled by phase-aligned and phase-inverted connections whose strength depends on their respective cell types. Connectivity is shown for one example instantiation of the network, with highlighted connections for a pair of opposing columns (at positions 0 and *π*). (B) Cell-type-level summary of connections in (A). (C) At each time point, the activity profiles of the arrays in the network are sinusoids of period 2*π*.

Here, *v*(*θ, t*) denotes the firing rate of the input neuron in column *θ* at time *t*. The phase of this input profile encodes the agent’s direction of movement arg(**v**) ∈ [0, 2*π*), and its amplitude encodes the agent’s translational speed ∥**v**(*t*)∥, scaled by a constant *k*_*V*_ that translates units of speed into firing rate. Because firing rates cannot be negative, the sinusoid includes a non-zero mean that ensures its trough is at zero, rather than a negative value. This velocity input is delivered in parallel to multiple columnar cell types, each of which weights it by a cell-type-specific connection strength *b*′_*i*_.

We then model the neural dynamics that result from this connectivity using a linear recurrent circuit model. In it, each columnar neuron acts as a leaky integrator of its phase-aligned, phase-inverting, and velocity inputs (Fig. 2A,B):

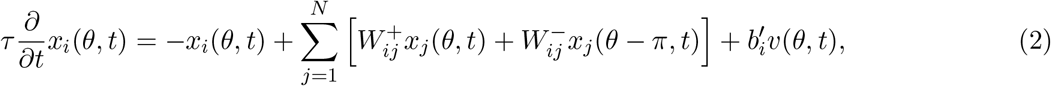

where *τ* is a neuronal time constant and *x*_*i*_(*θ, t*) denotes the firing rate at time *t* of the neuron of cell type *i* in column *θ*.

Due to the sinusoidal shape of the external input and the structure of the network’s dynamics, each cell type’s spatial activity profile will always be a sinusoid of period 2*π*, consistent with the kinds of spatial activity patterns observed experimentally. Thus, each activity profile is fully described by its first harmonic 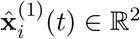 (which describes the sinusoid’s phase and amplitude) and its mean *µ*_*i*_(*t*) ∈ ℝ_+_ (Fig. 2C; Appendix A):

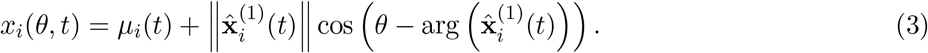

In the following section, we will illustrate how this network temporally processes the velocity input. Next, we will specify how network activity influences the agent’s velocity, thereby closing the loop.

### 2.2 Temporally filtering the velocity input

In this section, we will explore how the agent’s velocity shapes network activity. Recall that this network is composed of *N* arrays, and the activity of each array can be decomposed into two components: the first harmonic and the mean. These two components of network activity do not influence one another (Appendix A). Here, we will restrict our focus to the first harmonic (though see Appendix B for an overview of the mean component).

Let 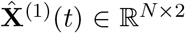 denote the state matrix whose rows contain the first harmonics of the *N* activity profiles. The dynamics of 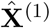 follow

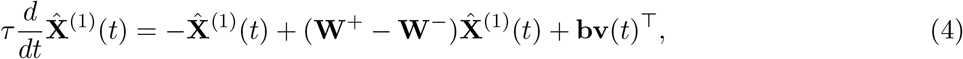

where **b** ∈ ℝ^*N*^ is a vector of input weights, with *b*_*i*_ = *k*_*V*_ *b*′_*i*_. Note that the recurrent couplings between the first harmonics depend on the differences between the phase-aligned and phase-inverted connection strengths through the coupling matrix **Ŵ** ^(1)^ = **W**^+^ − **W**^*−*^. This is because phase-inverting connections shift the influence of the presynaptic activity profile by *π* radians, which is equivalent to multiplying its first harmonic by −1 (Fig. 1E; [1]). Thus, even though all the single-neuron connections are excitatory, phase-inverting connections act as negative couplings between the first harmonics. Negative couplings promote repulsion between first harmonics, whereas positive couplings promote attraction (Fig. 3).

**Figure 3:**
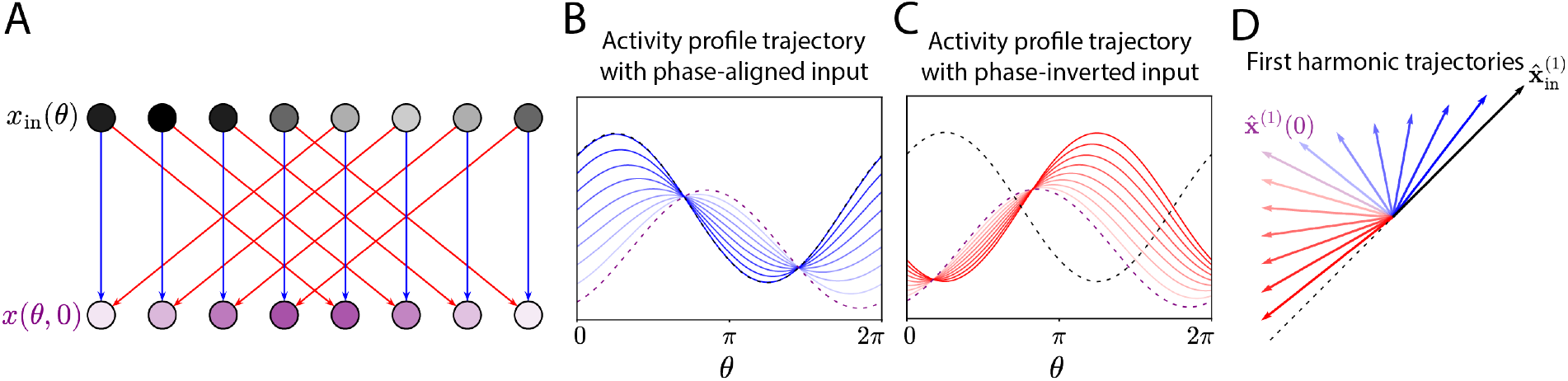
Dynamics of the first harmonic. (A) In this example, a neural array (purple) receives either phase-aligned (blue) or phase-inverted (red) connections from an input array (black) whose activity profile is constant over time. (B) The array’s activity profile is a time-varying sinusoidal pattern. When the input is phase-aligned, the activity profile flows from its initial state (purple dotted line) to that of the input array (black dotted line). (C) When the input is phase-inverted, the phase of the activity profile at steady state is opposite that of the input. (D) The nature of the phase shift determines whether the first harmonic of the activity profile is attracted towards or repelled by the first harmonic of the input.

To understand the computations being performed by this network, we next consider its responses to a time-varying input. Because the dynamics are linear and time-invariant, each array will temporally filter the network’s input using a kernel that depends on the network’s connectivity [25, 26]. The first harmonic of this input encodes the agent’s velocity vector. Consequently, the first harmonic of each activity profile will temporally filter the velocity history to compute a displacement-like quantity (Fig. 4A,B):

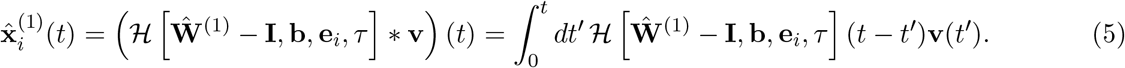

**Figure 4:**
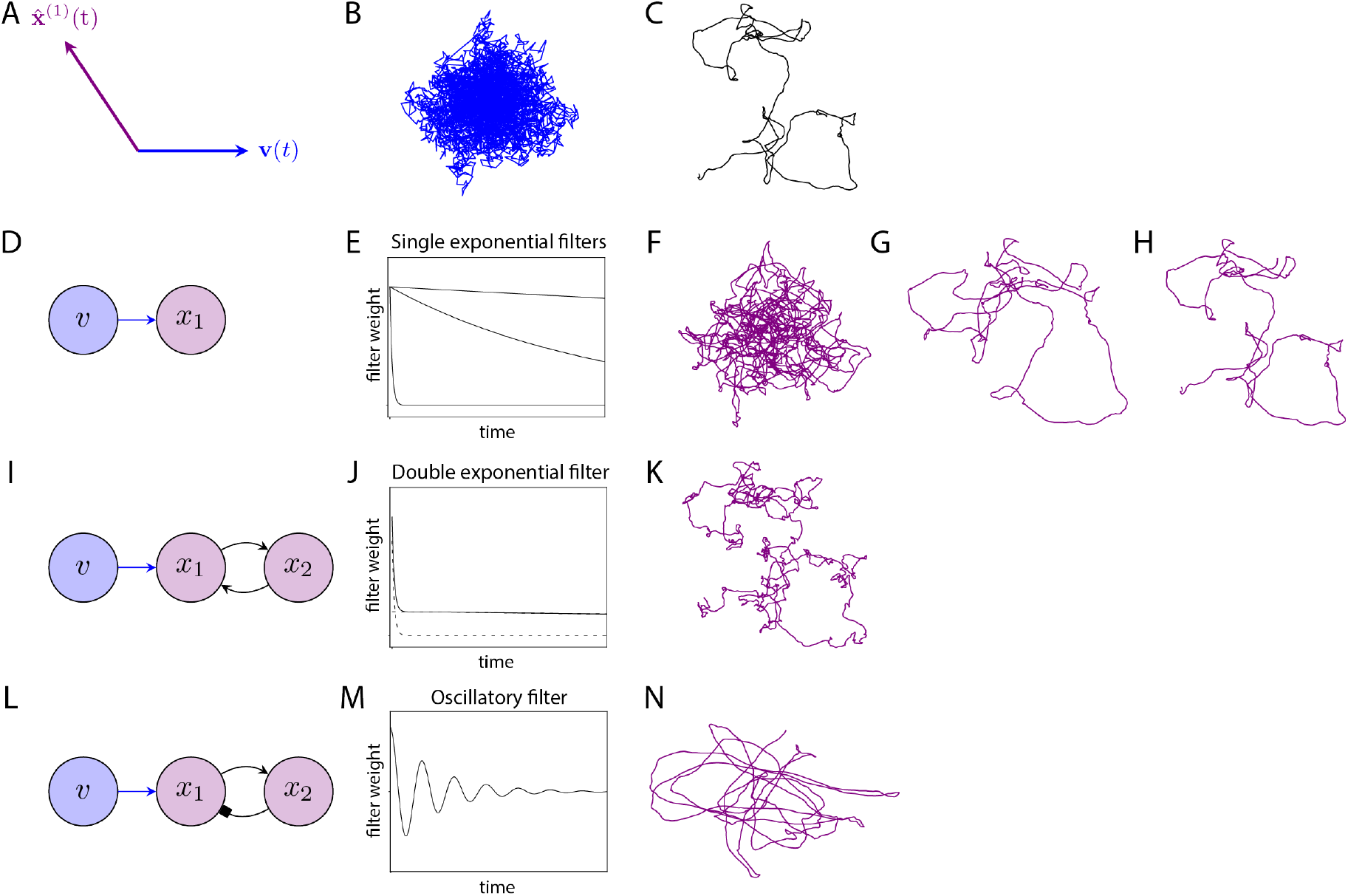
Temporally filtering the velocity input. (A) The first harmonic of each activity profile is a time-varying vector that depends on the sequence of velocity inputs and the network’s connectivity. (B) Example sequence of velocity vectors that acts as input in the examples that follow. (C) The associated positional trajectory. (D) A single self-coupled neural array integrating the velocity vector input. (E) Resulting temporal filters for different coupling strengths. (F-H) Resulting trajectories of the first harmonic of the neural array, ordered by increasing filter persistence. The most transient filter more closely tracks velocity, while the most persistent filter more closely tracks position. (I) A pair of rings coupled by phase-aligned connections. (J) The resulting filter is a double exponential. (K) This produces a first harmonic trajectory with structure on two distinct time scales. (L) A pair of asymmetrically coupled rings, in which one connection is phase-preserving and the other is phase-inverting. (M) The resulting filter is band-pass. (N) This generates a first harmonic trajectory that is oscillatory.

Above, *ℋ* […] is a functional that takes the parameters of a linear dynamical system (its internal couplings, input weights, readout weights, and time constant) and returns the temporal filter implemented by that system (Appendix C). The internal couplings here correspond to the network’s recurrent connectivity and leak, and they determine the filter’s characteristic time scales and oscillation frequencies. Meanwhile, the input and readout weights determine how strongly each of these components shapes a given feature of the system’s response to the input. In this case, the feature of interest is the first harmonic of the *i*th array, rather than a linear combination of multiple activity profiles, so the readout weights simply select that array using a one-hot vector **e**_*i*_ ∈ ℝ^*N*^.

When this filter is constant in time (e.g. *ℋ* […](*t*) = 1 for all *t*), the first harmonic tracks the agent’s positional displacement (Fig. 4C); this is classical path integration ([14]; see also [27–29]). However, the network can also realize more varied temporal filters, the shapes of which will depend on its connectivity (Appendix C). For example, a single-array network will act as a leaky integrator of the input, producing a smoothed estimate of recent displacement over the filter’s effective time window (Fig. 4D-H). In a multi-array network, each array can instead filter the input with a sum of exponentials, thereby mixing information about displacement over different time scales (Fig. 4I-K). Finally, a network with phase-inverted connections and oscillatory dynamics can also apply sinusoidal band-pass filters to the input (Fig. 4L-N). These networks generalize path integration to the wider variety of signal processing operations that can be realized by linear dynamical systems, enabling the network to extract information about the agent’s trajectory over various time scales and frequency bands.

### 2.3 Coupling behavior to neural activity through motor output

To close the loop, we need to specify how this network influences behavior. Recent experimental work on walking *Drosophila* has shown that the FB influences the animal’s heading and speed through two specialized populations of columnar output neurons: PFL2 and PFL3 [6, 15, 19]. Specifically, these output neurons continuously compare the fly’s current heading to a vector setpoint signal. When the heading is not aligned with the setpoint (e.g., when the fly turns away from the setpoint direction to inspect an object it encounters along its path, or to avoid an obstacle), these output neurons drive corrective turning and reduce forward speed, until the fly’s heading is once again aligned with the setpoint [15, 19]. The setpoint vector is encoded by a sinusoidal activity pattern that reflects the summed contributions of the FB columnar cell types that synapse onto PFL2 and PFL3 cells [6, 15, 18, 19].

Motivated by this experimental work, we model the setpoint signal as a weighted sum of activity profiles:

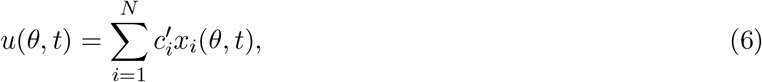

where 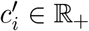 is the strength of the connections from neurons of cell type *i* to output neurons (PFL2 and PFL3 neurons) in the same column. Since this setpoint signal is a sum of sinusoids, it too is a sinusoid. Within our model, the first harmonic **û**^(1)^(*t*) of this sinusoid determines the instantaneous setpoint of the agent’s velocity vector:

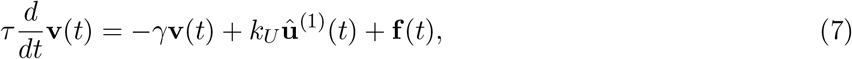

where *γ* is a drag-like coefficient that represents the agent’s intrinsic tendency to slow down, and *k*_*U*_ ∈ ℝ_+_ is a constant that specifies the magnitude of the locomotor drive that results from the setpoint signal. Equation 7 describes a control policy that continuously compares the velocity and setpoint vector and drives corrections when the two are misaligned. When rewritten in polar coordinates, this recapitulates the basic features of the heading and speed control performed by PFL2 and PFL3 cells (Appendix D). In addition, the velocity dynamics also depend on a forcing function **f** (*t*) ∈ ℝ^2^ that is independent of the FB-like network. This forcing function captures the time-varying influence of sensory stimuli and motor drives that influence the agent’s velocity vector through pathways that bypass the central complex. Finally, note that these velocity dynamics are linear. In Appendix G, we explore how nonlinearity in the transfer function that relates neural output to behavior affects the system’s dynamics.

### 2.4 Three descriptions of the closed-loop system

While neural activity influences the agent’s behavior through motor output, agent behavior also influences neural activity through the network’s velocity vector input (Fig. 5A). In this section, we consider three ways of reasoning about these reciprocal interactions.

**Figure 5:**
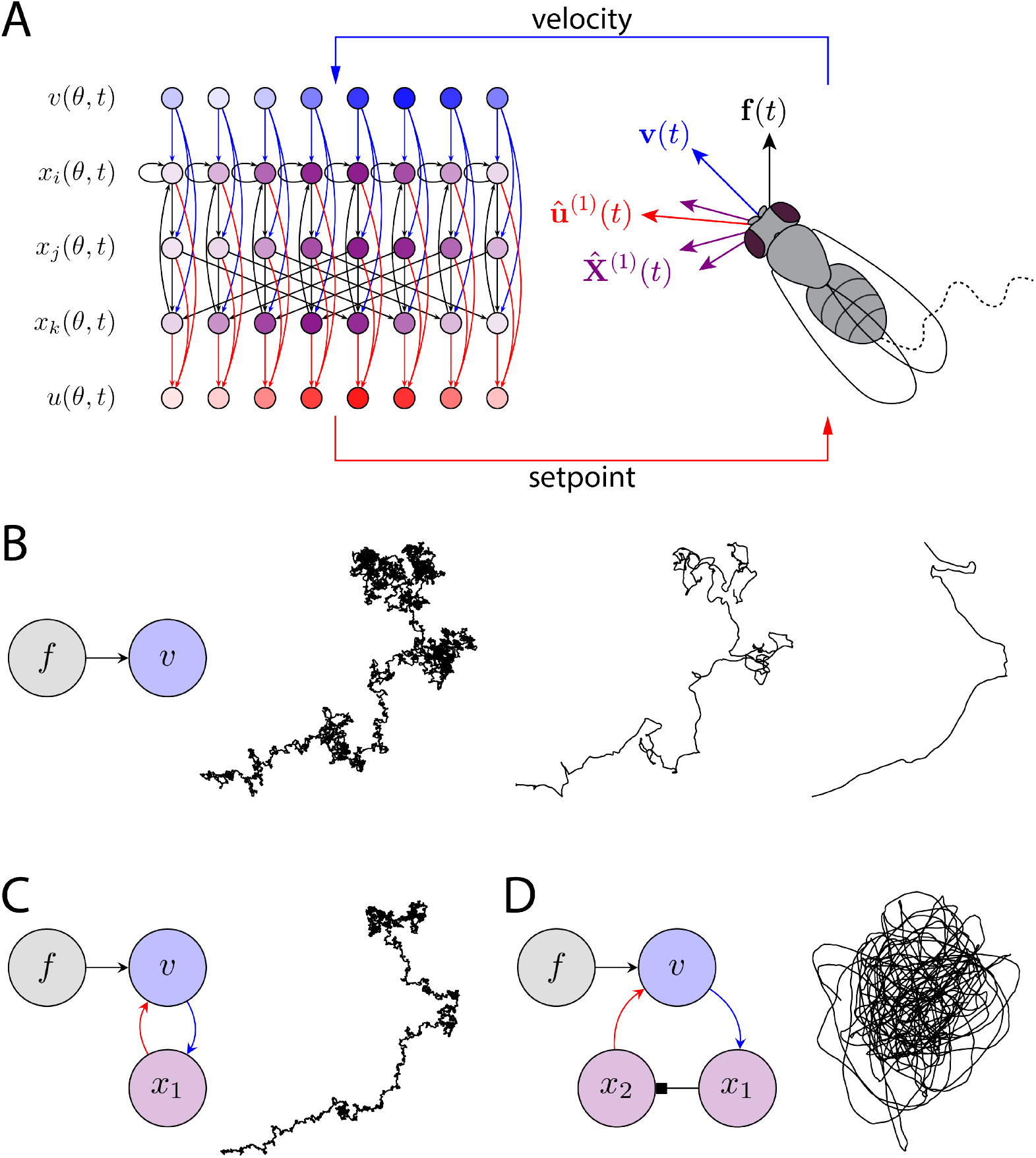
Shaping path structure. (A) The agent’s velocity vector and neural activity define a closed-loop system externally driven by the forcing function. (B) An agent with no arrays that is subject to random forcing produces a correlated random walk with a single correlation time scale that depends on *γ*. Decreasing *γ* increases path straightness. (C) The addition of one array produces a trajectory with structure on two time scales, resulting in small, diffusive excursions around a mostly straight path. (D) In response to the same forcing function, an agent with a phase-inverted connection produces an oscillatory trajectory. By inverting the phase of the velocity input in this manner, this network’s setpoint encourages periodic reversals.

First, we can describe the agent’s behavior as an autoregressive process that depends on the full history of past behavior. Here, we can eliminate neural variables altogether by rewriting the setpoint vector **û**^(1)^(*t*) as a filtered version of the velocity vector:

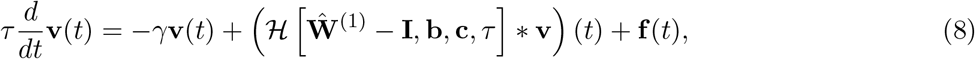

where **c** ∈ ℝ^*N*^ is a vector of output weights, with *c*_*i*_ = *k*_*U*_ *c*′_*i*_. The filter value *ℋ* […](*t* − *t*′) determines how strongly the current change in velocity is driven by the velocity at time *t*. Thus, the shape of the agent’s path depends on the shape of the filter, with more persistent filtering leading to more persistently correlated (or anti-correlated) behavior.

Second, we can describe behavior and neural activity as different dimensions of the same joint system:

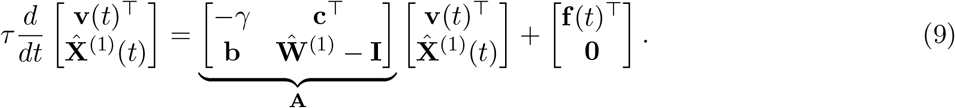

In this equation, the history dependence that was explicitly captured by the filter in equation 8 is instead summarized by the current activity pattern of each array. The dynamics of this system are set by the closed-loop coupling matrix **A** ∈ ℝ^(*N* +1)*×*(*N* +1)^. **A** jointly describes the network’s recurrent dynamics, the intrinsic (velocity-drag) dynamics of behavior, and the couplings between neural activity and behavior. From this perspective, the network’s connectivity influences behavior by influencing the properties of **A**, an idea known as pole placement in control theory [26].

Finally, we can describe the agent’s velocity as a filtered version of the motor drives captured by the forcing function:

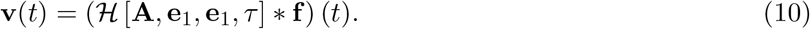

In this view, the dynamics of the joint system determine how the forcing function is transformed into a trajectory through the environment. As a result, the agent is not merely reactive, in that the forcing function alone does not dictate its instantaneous behavior. Rather, the network “observes” and “remembers” the agent’s past behavior, and this memory influences moment-to-moment responses to the forcing function.

In short, we can describe this system at several levels of abstraction. Across these levels, the FB network augments behavior through memory, determining how past behavior influences future behavior. In the sections that follow, we will illustrate how this can be useful.

### 2.5 Path control at multiple temporal and spatial scales

One potential use for this system is the control of the agent’s path structure, an important aspect of search behaviors like foraging. To illustrate this capacity, we will model the forcing function as an unstructured and random source of movement variability (white noise). This stochasticity might represent the random appearance of stimuli along the agent’s path, or random noise in the nervous system. Despite this unstructured forcing, the agent’s trajectory through the environment will possess a characteristic spatial and temporal structure that depends on the parameters of the closed-loop system. Thus, by adjusting the system’s couplings, the agent can tailor the structure of its exploratory behavior to its present needs and environment.

For example, an agent with no neural arrays will produce a trajectory with a single correlation time scale that depends on the drag coefficient *γ*. As in classic work on random walks [30, 31], varying this correlation time scale allows the agent to interpolate between highly diffusive movement that densely explores the area traversed and ballistic, straight-line movement that more efficiently moves between points (Fig. 5B). When neural arrays are introduced, the agent can generate richer paths that have structure on multiple spatial and temporal scales. For example, an agent with a single array can generate a trajectory that is relatively straight on long time scales but diffusive on short time scales, precisely because memory of recent travel direction allows it to resume course after an excursion (Fig. 5C). Finally, phase shifts enable an agent to generate oscillatory trajectories characterized by regular backtracking, which might be useful for more systematic local searches (Fig. 5D). These differences in path structure and their relation to the closed-loop system’s parameters are captured by summary statistics like the velocity autocovariance and mean squared displacement (Appendix E; equations 30, 31).

Note that in addition to altering the shape (e.g. straightness) of the trajectory, varying a given parameter in isolation can also alter the agent’s average speed. Independent control of path shape and speed can be achieved by a compensatory rescaling of the forcing function, which can be used to set the length scale of the trajectory while preserving its shape.

### 2.6 Stabilizing movement towards an intermittent cue

Another potential use for this system is the stabilization of movement guided by an intermittent cue, a previously proposed function of the FB [9, 17]. To illustrate this capacity, we augment the forcing function with a temporally sparse signal that biases the agent towards a fixed cue direction when that signal is present (Appendix F; Fig. 6A,B). For example, this signal might represent intermittent glimpses of a distant visual cue, or intermittent gusts of wind carrying an attractive odor. An agent that cannot stabilize movement in the cue direction will orient towards the signal when it is present but meander during the gaps between signals (Fig. 6C).

**Figure 6:**
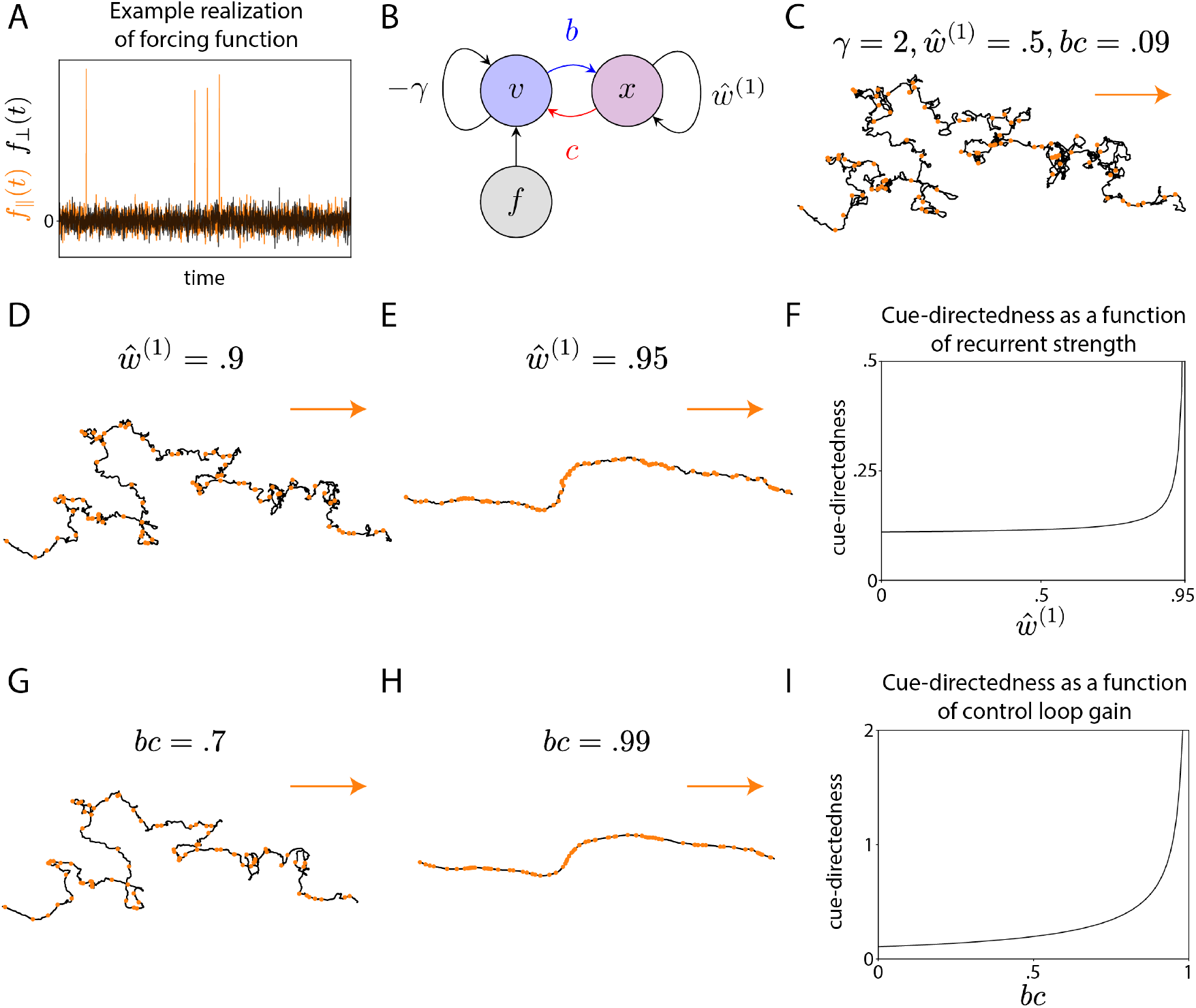
Stabilizing movement towards an intermittent cue. (A) The forcing function is a time-varying, two-dimensional signal that can be decomposed into components aligned (orange) and orthogonal (black) to the cue direction. It consists of a directionally-unbiased component that continuously injects variability into the agent’s behavior, and a cue-directed component that is intermittently present. (B) The properties of the closed-loop system determine the trajectory that results from this forcing. In this example, we consider a network with a single array. (C) Example positional trajectory. Orange dots correspond to moments when the intermittent cue is present. This cue transiently biases the agent in the cue direction, which points to the right. Here, the couplings are relatively weak, so the agent’s trajectory is meandering. (D,E) Example trajectories with the same forcing function but stronger recurrent connectivity, corresponding to a longer neural integration time scale. (F) The resulting path becomes more cue-directed (ratio of movement aligned and orthogonal to the cue direction; Appendix F) as the neural recurrence increases. (G,H) Example trajectories with higher control loop gain, which represents the strength of the network’s velocity input and/or steering output (Appendix F). (I) The path becomes more cue-directed as the loop gain increases.

In contrast, an agent whose behavior integrates the forcing function over a longer time scale will more persistently move in the cue direction during gaps in the signal (Fig. 6D-I). The specific time scale realized by the agent’s behavior will depend on the dynamics of the closed-loop system, and it can be manipulated in two ways. First, strengthening the network’s recurrent connections increases the time scale over which the setpoint signal integrates the agent’s velocity and, by extension, the forcing function. This enables the agent to accumulate evidence of the cue’s salience and amplify biases in the forcing function over longer time scales (Fig. 6D-F). Second, we can increase the gain of the control loop (determined by the strength *b* of the network’s velocity input and the strength *c* of the network’s motor output), which enables the agent to more strongly correct moment-to-moment differences between its velocity and setpoint. Thus, even if the setpoint signal itself only integrates the velocity over a small window, the FB network can increase persistence in the cue direction by more aggressively combating drift in its heading direction (Fig. 6G-I). In either case, by tuning the dynamics of the closed-loop, the agent can balance random exploration and exploitation of the intermittent cue.

## 3 Discussion

In this study, we have shown how an FB-like network can control the spatial and temporal structure of an agent’s paths through its environment. This recurrent network temporally filters a velocity vector input, and it uses this filtered vector as the setpoint of a controller that continuously adjusts the agent’s velocity. Importantly, the agent’s velocity is also influenced by lower-level sensorimotor pathways. These lower-level pathways allow the agent to react to local environmental features and inject variability into its behavior, while the higher-level recurrent network maintains a memory of past movement and governs the overall path structure.

This path control perspective may clarify several otherwise puzzling features of the FB, such as the multitude of pathways downstream of the velocity input, the prevalence of leaky integrator dynamics, and the circular relationship between the setpoint signal and the animal’s velocity vector. Our model suggests that the multitude of columnar cell types downstream of the velocity input may enable the FB to extract a variety of temporal or spectral features from the velocity trajectory. Meanwhile, using a combination of these features as a time-varying velocity setpoint may allow the FB to influence path structure at multiple spatial and temporal scales. This work thus builds on previous models of path integration in the FB [6, 8, 9, 14, 32] by exploring circuit implementations of more varied temporal filters.

This work also leverages the idea that sinusoidal activity patterns can represent two-dimensional vectors [1, 14]. In previous work, these activity patterns have encoded variables like an agent’s velocity [4, 5], positional displacement [6, 8, 9, 14], and the direction and speed of the wind [33]. In contrast, individual cell types in our model do not explicitly encode a specific environmental variable. Instead, each array simply applies a cell-type-specific linear filter to the velocity input, and this filtering results in activity patterns that help generate behavior. This distinction between representation and generation loosely echoes discussions of neural dynamics in areas like motor cortex [34, 35].

In addition to its first harmonic (sinusoidal component), each activity profile also has a time-varying mean activity level. Within the FB literature, the mean has received less attention, in part because it lacks a clear interpretation that is analogous to the first harmonic’s interpretation as a vector. In Appendix B, we analyze the dynamics of mean activity in our model. There, we show that much like the first harmonics temporally filter the agent’s velocity vector, the means temporally filter the agent’s translational speed. Thus, much like the first harmonic acts as a displacement-like vector variable, the mean acts as a path-length-like scalar variable. We also note that the first harmonic and mean differentially interact with phase-inverting connections, which negatively couple the first harmonics of the activity profiles but positively couple their means. Finally, while the mean plays no role in our model’s simple control loop, this simplification is unlikely to hold in practice: optogenetically increasing the mean activity of FB output neurons can influence locomotion [19], and several cell types downstream of the FB are well-poised to read out the mean activity of particular FB columnar cell types [3].

Another idealized feature of the behavioral dynamics (equation 7) we analyzed in the main text was their linearity. An unrealistic consequence of this linearity is that the agent’s speed is unbounded. To address this, in Appendix G, we augment the behavioral dynamics with a saturating nonlinearity that prevents the agent’s speed from exceeding some maximum value. There, we note that the introduction of this simple nonlinearity can result in an interesting emergent phenomenon: self-sustaining neural activity and behavior. In this regime, the neural circuit’s persistent activity is sustained by the sensory input that results from its own motor output. This mirrors, and to some degree contrasts with, mechanisms of persistent activity that instead rely on strong recurrent interactions between neurons [36, 37]. It also functions as a simple example of structure jointly emerging from the dynamics of “the body” (the agent’s behavior) and the nervous system, a principle commonly stressed in embodied cognition [38–40].

Regardless of the precise mapping from neural activity to behavior, the general function of this model is to control path structure. The resulting paths can combine persistent, meandering, and oscillatory movement, and their exact structure depends on the model’s coupling parameters. Interestingly, these paths are examples of correlated random walks. Correlated random walks are commonly used descriptive models in the fields of animal movement and computational ethology [30, 41, 42]. Here, we illustrate one way that a correlated random walk with a given statistical structure can directly arise from an underlying network model.

Correlated random walks with different structures can be useful in different contexts of foraging and search. According to foraging theory, organisms need to employ a contextually appropriate mix of persistent and meandering movement to search an environment [43, 44]. Meandering allows an organism to densely explore a rich environment, while straight movement allows it to rapidly leave a barren one. The right strategy depends on the organism’s environment and internal state. This contextual information may reach the FB via tangential cells [3]. The arbors of individual tangential cells innervate all the columns of a given FB layer. This morphology makes these cells well-poised to modulate columnar activity dynamics and thereby tailor the organism’s trajectories to its present circumstances [17, 45].

Of course, organisms that lack a nervous system can still forage. For example, bacteria employ a run-and-tumble foraging strategy, in which straight runs are punctuated by random turns whose frequency depends on environmental cues [46]. Whereas idealized models of run-and-tumble foraging are memoryless, our network model temporally filters the velocity input to maintain a memory of past motion. As a result, this network can influence path structure on a variety of time scales and spatial scales. For example, the path can be straight on long time scales but meandering on short ones, precisely because memory allows it to resume course after an excursion.

Another function of this network is to stabilize useful actions, based on accumulated evidence in favor of those actions [9, 17, 47]. This capacity is useful because natural sensory signals can be discontinuous in time. For example, an attractive visual object can be intermittently visible, or a breeze carrying a food odor can be intermittently sensed. Within this model, orientation towards such cues is initiated by lower-level sensorimotor systems that are independent of the FB-like network. These lower-level systems are represented by the forcing function. When the forcing function exhibits a directional bias due to the intermittent signal, the network can stabilize movement in this direction regardless of the signal’s modality or valence. Thus, the system can sustain actions initiated by more “distractable” low-level pathways, a capacity loosely reminiscent of executive function. Interestingly, *Drosophila* will navigate toward remembered cue directions, but genetic disruptions of the compass system disrupt this ability [48].

This dependence on the central complex is consistent with the idea that persistent navigation towards a remembered cue emerges from FB network dynamics.

In many ways, the FB exhibits the rudiments of cognition: it contains allocentric, almost symbolic representations of certain variables; can manipulate these representations using phasor-based arithmetic operations; exhibits working memory; accumulates evidence over time; and enables persistent action that is robust to distractors. Interestingly, analogous high-level functions are sometimes attributed to brain regions like the prefrontal cortex [49]. It is often natural to view such regions as the apex of a control hierarchy, issuing abstract commands that low-level systems translate into motor output. In this closed-loop model, however, the low-level pathways represented by the forcing function are the real engine of behavior, while the high-level network mainly influences the time scales over which the behaviors they initiate persist. Recent findings suggest that the prefrontal cortex may function similarly during self-directed exploratory behavior [50].

Within this model, there is also no direct neural communication between these low-level pathways and the network. Instead, the network simply observes the influence of these pathways on behavior itself, and it bases its setpoint for future behavior on current behavior. The idea that neural controllers operating in parallel might interact through their mutually observed effects on behavior—rather than internally agreeing on some coherent course of action prior to carrying out—has some precedent in cognitive science and robotics [51, 52].

Unlike other recent models of agents controlled by recurrent neural networks [53, 54], we did not train this network to perform a task. Instead, we aimed to understand the neural dynamics and behavior that could emerge from observed patterns of neural connectivity in the insect central complex. Closed-loop models of network-driven behavior also have precedents in the motor control literature. Here, neuromechanical models have explored how biomechanics and neural circuitry jointly produce locomotion [55–58]. Our work extends closed-loop modeling from the sensorimotor periphery (e.g. the spinal cord and brainstem) to a higher-order structure, the central complex. Although the central complex is deep in the brain, the input and output pathways that link its activity to behavior are known explicitly and thus well-constrained.

In our model, the network’s structure resulted in a low-dimensional space of neural activity patterns, defined by the first harmonic of each activity profile. The network’s sensory input and motor output could both be defined with respect to this space (rather than single neuron activities), and the simplicity of the resulting interactions enabled us to directly characterize the joint system consisting of both neural and behavioral variables. This approach therefore connects models of neural computation based on the population dynamics of recurrent circuits [59, 60] to models of feedback control based on reciprocal interactions between the brain, body, and external world [38–40, 61].

## Acknowledgments

Many thanks to colleagues in the Wilson and Pehlevan Labs for their feedback on this work, and to Caleb Weinreb and Jacob Zavatone-Veth for their comments on an earlier version of this manuscript. During the preparation of this manuscript, M.A.M.O. used GPT-5 to generate code and proofread the text, via a ChatGPT Edu license provided by the Harvard Kenneth C. Griffin Graduate School of Arts and Sciences.

C.P. is supported by an NSF CAREER Award (IIS-2239780) and the Simons Collaboration on the Physics of Learning and Neural Computation. This work has been made possible in part by a gift from the Chan Zuckerberg Initiative Foundation to establish the Kempner Institute for the Study of Natural and Artificial Intelligence. This work was also supported by NIH grant R01NS129647 to R.I.W. R.I.W. is an HHMI Investigator.

## Appendix

### A Dynamics of the Fourier coefficients

In this section, we derive the governing equations for the Fourier coefficients (first harmonic and mean) of the activity profiles. The model in the main text is an example of a spatially organized network with multiple cell types and translationally invariant connections [62–64]. This connectivity is organized around an angular variable, more specifically making the model a ring network [29, 36, 65, 66]. In the linear case, the dynamics of such systems can be expressed as follows:

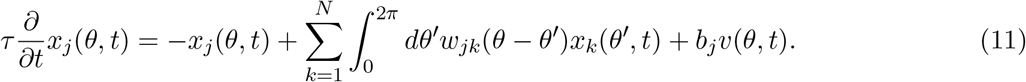

Here, *v*(*θ, t*) is some external input and *w*_*jk*_(*·*) is a connectivity kernel (periodic over 2*π*) that relates the strength of a connection to the cell types and relative positions of the two neurons. For simplicity, we model the network as a continuum, though translating to the discrete case is straightforward (see [29]).

As is standard in the study of such networks, we can use the Fourier transform to express the shape of each population’s activity profile in the frequency domain. Let 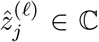 denote the *ℓ*th (with *ℓ* ∈ *N*) unnormalized Fourier coefficient of the *j* th activity profile:

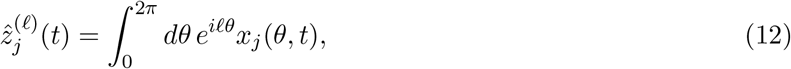

where 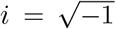 denotes the imaginary unit (used elsewhere in this manuscript as a discrete indexing variable). Similarly, let 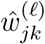 and 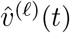 denote the *ℓ*th Fourier coefficients of the connection kernels and input pattern. The dynamics of 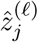 follow

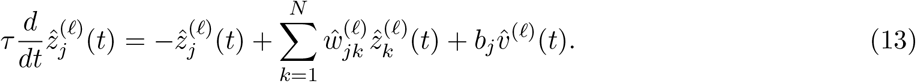

Note that 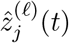 interacts with the *ℓ*th Fourier coefficients of the other activity profiles and the input but is completely uncoupled from Fourier coefficients corresponding to other frequencies (*m*≠ *ℓ*). Each Fourier mode thus defines an independent dynamical system driven by the corresponding Fourier coefficient of the input and coupled through the corresponding Fourier coefficients of the connectivity kernels:

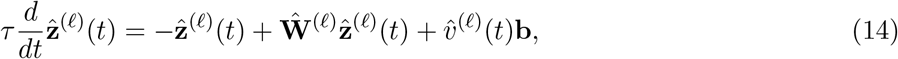

where 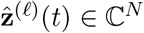 is the population vector with entries 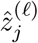 and **Ŵ** ^(*ℓ*)^ ∈ C^*N×N*^ is the coupling matrix with entries 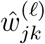 (see [29] and [64] for analogous modal reductions).

Because the external input in the main text is a simple sinusoid lacking any high frequency content 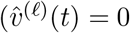 for all *ℓ >* 1), its activity profiles will not exhibit high frequency content unless it is present at initialization. In addition, assuming a given mode’s dynamics are stable (Re *λ*_*j*_ **Ŵ** ^(*ℓ*)^ *<* 1 for all *j*), any high frequency content present at initialization will dissipate. In any case, we assume an initial state of zero (*x*_*j*_(*θ*, 0) = 0) throughout, thus guaranteeing that the activity profiles are always simple sinusoids and reducing the dynamics to modes 0 and 1. In the main text, for clarity, we expressed these modes using real (rather than complex) numbers and rescaled them so that their magnitudes directly equal the associated features (mean and amplitude) of the activity profile:

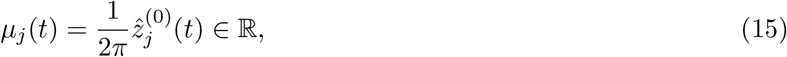

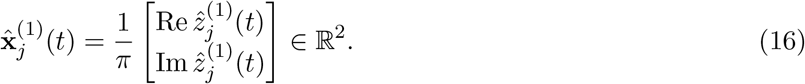

Finally, to obtain the governing equations for these variables, we consider the specific connectivity kernel used in the main text. This connectivity kernel is a weighted sum of two delta functions, respectively corresponding to the phase-preserving and phase-inverting connections:

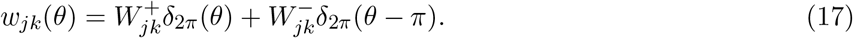

The Fourier coefficients of this kernel depend on the parity of the mode:

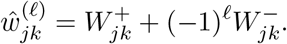

This parity dependence reflects the fact that a phase shift of *π* leaves modes of even wavenumber unchanged while multiplying modes of odd wavenumber by −1. Consequently, phase-inverting connections act as negative, “effectively inhibitory” couplings among the first (and other odd-numbered) harmonics of the activity profiles, even though all the units in the network are excitatory.

### B Dynamics of the mean

The activity of each neural array in our model can be decomposed into two components: the first harmonic and the mean. In this section, we consider the dynamics of the latter. Let ***µ*** ∈ ℝ^*N*^ denote the mean activity in each array. The dynamics of ***µ*** follow

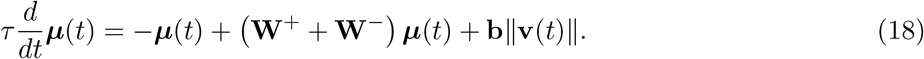

Whereas the first harmonic of the network’s input encodes the agent’s velocity vector, the mean of the input encodes the agent’s translational speed (due to the non-zero offset in equation 1). As a result, whereas the first harmonic computes a displacement-like quantity by temporally filtering its velocity vector input, the mean computes a path-length-like quantity by temporally filtering its speed input. In addition, note that phase-inverting connections positively couple the means but negatively couple the first harmonics. Due to this coupling difference, the two components will generally apply different filters to their respective inputs when the network contains phase-inverting connections.

Finally, since the network’s connections and input are non-negative, single-neuron activity must also be non-negative. As a result, the mean of each array’s activity profile must always equal or exceed the amplitude of its first harmonic. Since the mean bounds the magnitude of the first harmonic, the stability of the network solely depends on the stability of the mean dynamics, with stability condition *λ*_max_(**W**^+^ + **W**^*−*^) *<* 1.

### C Linear filtering

Many of the results in the main text rely on the fact that time-invariant linear dynamical systems act as temporal filters of their inputs. In this section, we review the relevant mathematics, which constitute standard linear systems theory (see e.g. [25, 26]). Consider a system of the form

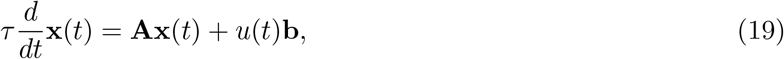

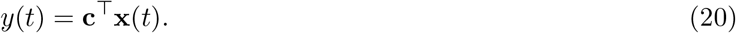

Here **x**(*t*) ∈ ℝ^*N*^ is the system state, *u*(*t*) ∈ ℝ is some time-varying input to the system, *y*(*t*) ∈ ℝ is some observed variable, **A** ∈ ℝ^*N×N*^ is a matrix of recurrent couplings, **b** ∈ ℝ^*N*^ is a vector of input weights, and **c** ∈ ℝ^*N*^ is a vector of the readout weights. Again, we will assume an initial condition of **x**(0) = **0**.

We can express *y*(*t*) as some convolution of *u*(*t*) with a kernel that depends on the system’s coupling parameters:

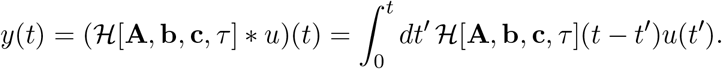

Here, *ℋ* is a functional that takes in the system parameters and returns the associated temporal kernel:

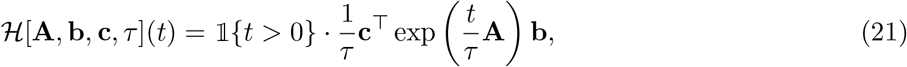

where the coefficient 1{*t >* 0} ensures that the kernel is causal (equals 0 for *t <* 0). In general, such a filter can be expressed as a sum of up to *N* terms, each consisting of a polynomial times an exponential. Thus, for *t >* 0:

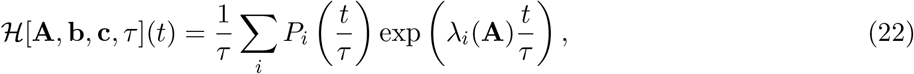

where 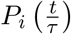 is a polynomial function of time and *λ*_*i*_(**A**) ∈ C denotes the *i*th eigenvalue of **A**.

If the matrix **A** is diagonalizable, then each 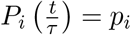 will be a constant and the filter will simply be a weighted sum of exponentials. Non-diagonalizable systems with nontrivial polynomial terms can exhibit interesting filtering properties that have been explored elsewhere [28, 67].

The behavior of exponential terms will in turn depend on the eigenvalues of **A**. Real-valued eigenvalues will contribute exponential decay terms that (leakily) integrate the input with an effective time constant of 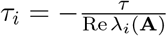. Complex-valued modes will come in conjugate pairs, resulting in exponentially damped (or driven) oscillatory terms of the form

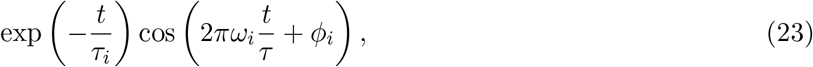

where *ω*_*i*_ ∈ ℝ determines the oscillation’s frequency and *ϕ*_*i*_ ∈ [0, 2*π*) determines its phase. These oscillatory terms can realize more complex (e.g. band-pass) filtering properties.

With this background in mind, we can summarize many of the variables in the main text as temporally filtered responses to external inputs:

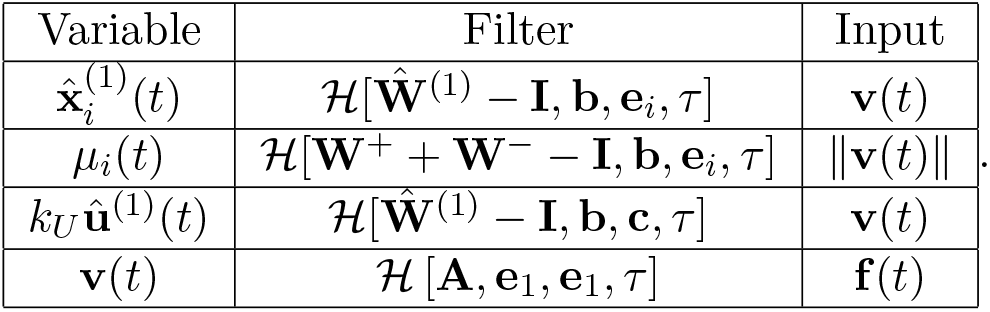

Above, **e**_*i*_ denotes the *i*th standard basis vector (the vector with entries *δ*_*ij*_).

### D Polar coordinate description

In this section, we reexpress the dynamics of the closed-loop system in polar coordinates. This change of coordinates yields governing equations for the agent’s heading and forward speed, thereby making connections to the literature on central complex output circuits more apparent. It also yields governing equations for the activity profile phases and amplitudes, thereby providing a different perspective on the neural dynamics.

To start, we will define polar representations for each of the variables. Let 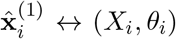 where *X*_*i*_ denotes the amplitude of the *i*th activity profile and *θ*_*i*_ denotes its phase; **û**^(1)^ ↔ (*U, ψ*), where *U* denotes the amplitude of the setpoint signal and *ψ* denotes its phase; and **v** ↔ (*V, ϕ*), where *V* denotes the agent’s forward speed and *ϕ* denotes its heading. Note that “heading” simply refers to the direction of the velocity vector in this model. Since the agent is modeled as a particle without an intrinsic axis of orientation, the model does not distinguish between heading and direction of travel.

When equation 7 is rewritten in polar coordinates, the resulting heading dynamics recapitulate the basic features of the steering control loop implemented by PFL3 cells [6, 15, 19]:

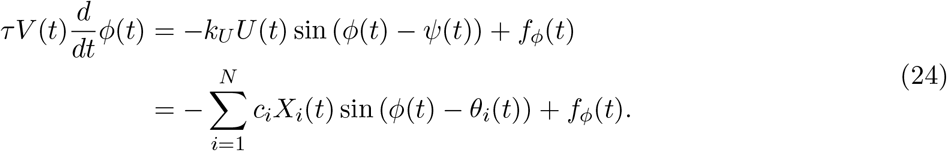

This equation continuously compares the agent’s current heading *ϕ*(*t*) to the setpoint phase *ψ*(*t*), yielding an “alignment error” sin(*ϕ*(*t*) − *ψ*(*t*)). The resulting heading update is then proportional to the negative error, which causes the agent to turn towards the setpoint phase. In our model, the gain of this turning drive scales with *U* (*t*), the amplitude of the setpoint signal. Additionally, as the second line of equation 24 shows, we can also decompose the full turning output into a weighted sum of analogous contributions from the *N* arrays. From this latter perspective, each array steers the agent’s heading towards its activity phase *θ*_*j*_(*t*). The strength of this pull depends on the amplitude *X*_*j*_(*t*) and the output weight *c*_*j*_. In our model, turning gain is also inversely proportional to forward speed *V* (*t*). Finally, *f*_*ϕ*_(*t*) ∈ ℝ denotes the effect of the forcing function orthogonal to the current heading.

The radial component of the polar dynamics relates the agent’s forward speed to the degree of alignment between the heading and setpoint, as well as the amplitude of the setpoint signal. There are populations of output neurons—namely, PFL2 cells [19]—well-poised to implement such a speed controller. More formally,

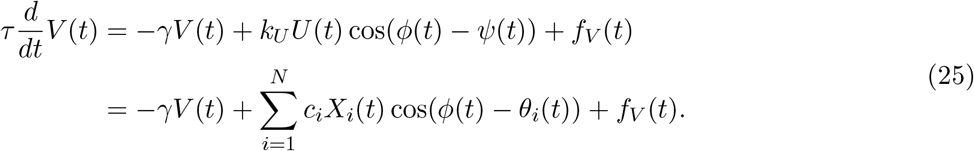

This equation consists of a drag-like term that drives forward speed to 0 in the absence of input. The neural drive that combats this tendency is proportional to cos(*ψ*(*t*) − *ϕ*(*t*)) ∈ [−1, 1], which quantifies the degree of alignment between the setpoint signal and heading. Ignoring the forcing function and treating the other state variables as fixed, this equation will drive forward speed to a maximal value 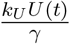 when heading and setpoint are perfectly aligned, a speed of 0 when the two are orthogonal, and a speed of – 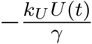 when the two are oppositely oriented. We can view moments of transition from positive to negative *V* as instantaneous inversions of the heading rather than backwards movement, since the points (−*V, ϕ*) and (+*V, ϕ* + *π*) are mathematically equivalent in polar coordinates. Like the turning dynamics, the forward speed dynamics can also be decomposed into a linear combination of analogous per-array contributions (second line of equation 25). In addition, *f*_*V*_ (*t*) ∈ ℝ denotes the effect of the forcing function aligned with the current heading.

Finally, to understand the joint dynamics of neural activity and behavior, we rewrite the neural dynamics along the first harmonic in polar coordinates. The phase dynamics are given by

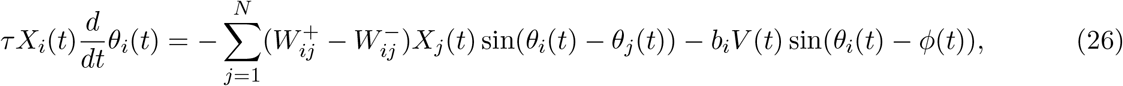

which is analogous to the governing equation that relates neural activity to changes in heading. Similarly, the amplitude dynamics are given by

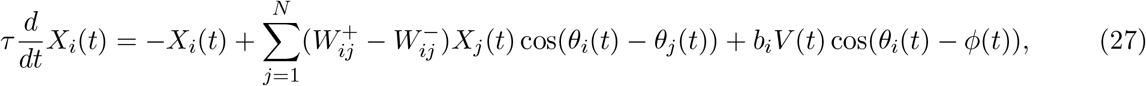

which is analogous to the equation relating neural activity to changes in forward speed. In summary, the phases of the neural activity bumps and the agent’s heading define a set of angular variables coupled by alignment (and amplitude-dependent) interactions, somewhat akin to the synchronizing forces in a Kuramoto model [68].

### E Statistical structure of random walks

To illustrate how the network’s couplings can shape the agent’s trajectories through its environment, we model the forcing function as a minimally structured source of random variability:

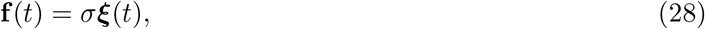

where ***ξ***(*t*) ∈ ℝ^2^ is a white noise process with ⟨***ξ***(*t*)⟩ = 0 and ⟨***ξ***(*t*)***ξ***(*t*′)^*⊤*^⟩ = *δ*(*t* − *t*′)**I**, where ⟨·⟩ denotes an expectation over noise realizations. In addition, *σ* ∈ ℝ_+_ is a constant that sets the noise strength. With this stochastic forcing, the agent will perform a correlated random walk through its environment [30, 31]. The overall trajectory will reflect a weighted sum of the *N* + 1 modes present in the joint system. For concreteness, we will assume the closed-loop coupling matrix **A** is diagonalizable and its eigenvalues are real, so that the temporal filter realized by the agent’s velocity is a sum of exponentials of the form

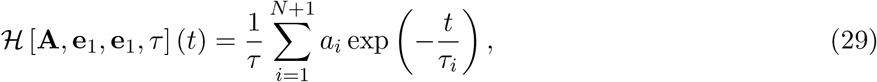

where *a*_*i*_ ∈ ℝ and 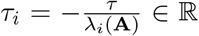 are the coefficient and effective time constant associated with the *i*th mode of the filter. We further assume that the joint system is stable (*λ*_*i*_(**A**) *<* 0 for all *i*).

With such a filter, the random walk will exhibit a multiplicity of time scales that are evident in its autocovariance. The autocovariance relates the expected similarity (inner product) of the velocity vector at two time points to the duration Δ ≥ 0 separating them. In this case, the autocovariance of the agent’s random walk will contain up to *N* + 1 modes that are inherited from the filter:

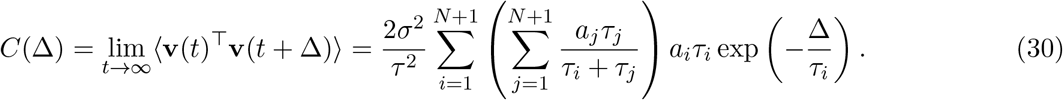

Each term in this weighted sum contributes a correlation time scale of *τ*_*i*_. Terms with positive coefficients promote persistent movement over lags Δ *< τ*_*i*_, while terms with negative coefficients promote reversals over the same time scale. The overall path structure will reflect the combined contributions of each mode. The filter’s time scales are also present in the agent’s mean squared displacement (MSD), which characterizes how the agent’s distance from the starting point of an interval scales with the interval’s duration:

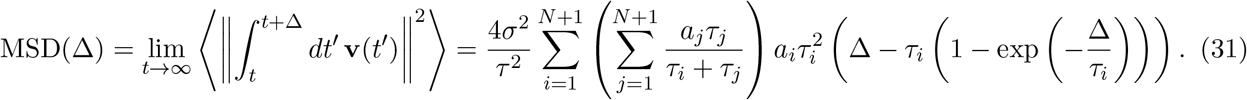

Finally, if the joint system has complex modes, the filter will contain exponentially damped oscillatory terms. Such modes will encourage periodic backtracking in the agent’s trajectory and contribute oscillatory terms with the same frequency and decay rate to the autocovariance.

### F Navigation towards an intermittent signal

To model navigation towards an intermittent sensory cue, we augment the forcing function with a temporally intermittent signal—mathematically equivalent to “shot noise” [31, 69]—that biases the agent towards some cue direction **z** ∈ ℝ^2^ (with ∥**z**∥ = 1):

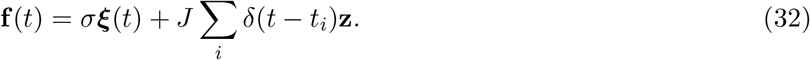

In this equation, {*t*_*i*_} defines a set of observation times sampled from a Poisson process with rate parameter *v*, and *J* ∈ ℝ_+_ parameterizes the strength of each signal’s immediate effect on behavior. To characterize the agent’s propensity to move in the cue direction, we decompose its velocity into components aligned with and orthogonal to **z**, denoted *v*_*∥*_(*t*) and *v*_*⊥*_(*t*). Let 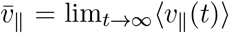 denote the mean cue-aligned velocity and 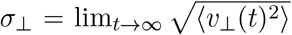 denote the standard deviation of the cue-orthogonal velocity. The Ratio 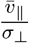 is then a natural measure of cue-directedness, somewhat akin to a signal-to-noise ratio.

For a system with closed-loop coupling matrix **A**, the cue-directedness is given by

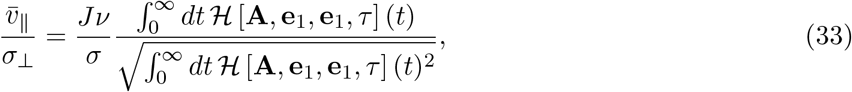

where *ℋ* [**A, e**_1_, **e**_1_, *τ*] is the kernel with which the agent’s velocity filters the forcing function (equation 10). This equation consists of two factors. The first factor captures the contribution of the forcing function itself. It is directly proportional to the cue frequency and strength and inversely proportional to the magnitude of the isotropic noise. The second factor captures the closed-loop interactions between the network and behavior. Its numerator corresponds to the DC gain with which the system amplifies signal in the forcing function, while its denominator captures the variance generated by filtering the white noise.

In Figure 6, to explore how the network’s dynamics can influence the agent’s behavior, we varied the parameters in a network consisting of a single array (*N* = 1) and measured the cue-directedness of the resulting trajectories. More formally,

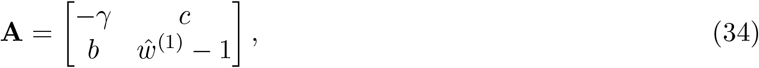

where *γ, b, c >* 0, ***Ŵ***^(1)^ *<* 1, and *γ*(1 − ***Ŵ***^(1)^) *> bc* (which ensures stability). We then showed that given the same forcing, the cue-directedness of the resulting path increases as ***Ŵ***^(1)^ (which sets the neural integration time scale) and *bc* (which sets the gain of the control loop) increase. Thus, varying both the integration time scale of the network and the gain of the closed loop can enable the agent to exploit this intermittent source of information and stabilize coherent movement in the cue direction.

### G Dynamics with velocity nonlinearities

One unrealistic feature of the linear velocity dynamics defined by equation 7 is that the agent’s speed is unbounded. This can be addressed by augmenting the velocity dynamics with a saturating nonlinearity that ensures the agent stays below some maximum speed:

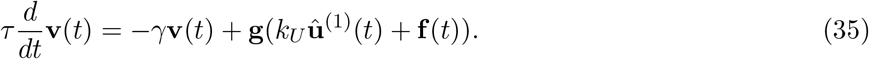

Here **g** : ℝ^2^ → ℝ^2^ is a vector-valued function that rescales its input vector by applying a saturating (tanh) nonlinearity to its magnitude while preserving its direction:

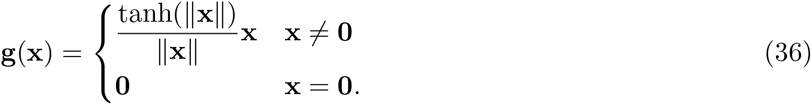

Note that this nonlinearity solely alters the dynamics of the velocity vector. The neural dynamics remain linear, following equations 2 and 4. When this speed nonlinearity is introduced, the basic phenomena noted in the preceding sections still hold: increasing the strength of the neural recurrence and the gain of the control loop both increase the straightness of the simulated trajectories.

However, new phenomena also emerge. For example, in the absence of forcing (**f** (*t*) = **0**), a linear agent (with stable closed-loop couplings) will remain stationary, since its dynamics are dissipative and its behavior is input-driven. The same is true of a nonlinear agent with weak couplings. However, as the nonlinear agent’s coupling strengths increase, it can undergo a transition from this quiescent state to an active state in which neural activity and behavior are self-sustaining. This transition can be observed in the single array setup defined by equation 34. To analyze it, we can linearize the dynamics of the agent’s translational speed *V* (*t*) and neural amplitude *X*_1_(*t*) (derived from equations 25 and 27) around the stationary fixed point *V* (*t*) = *X*_1_(*t*) = 0:

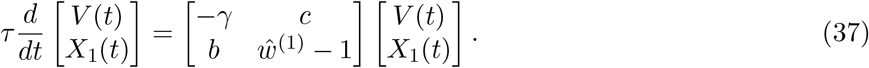

**Figure S1:**
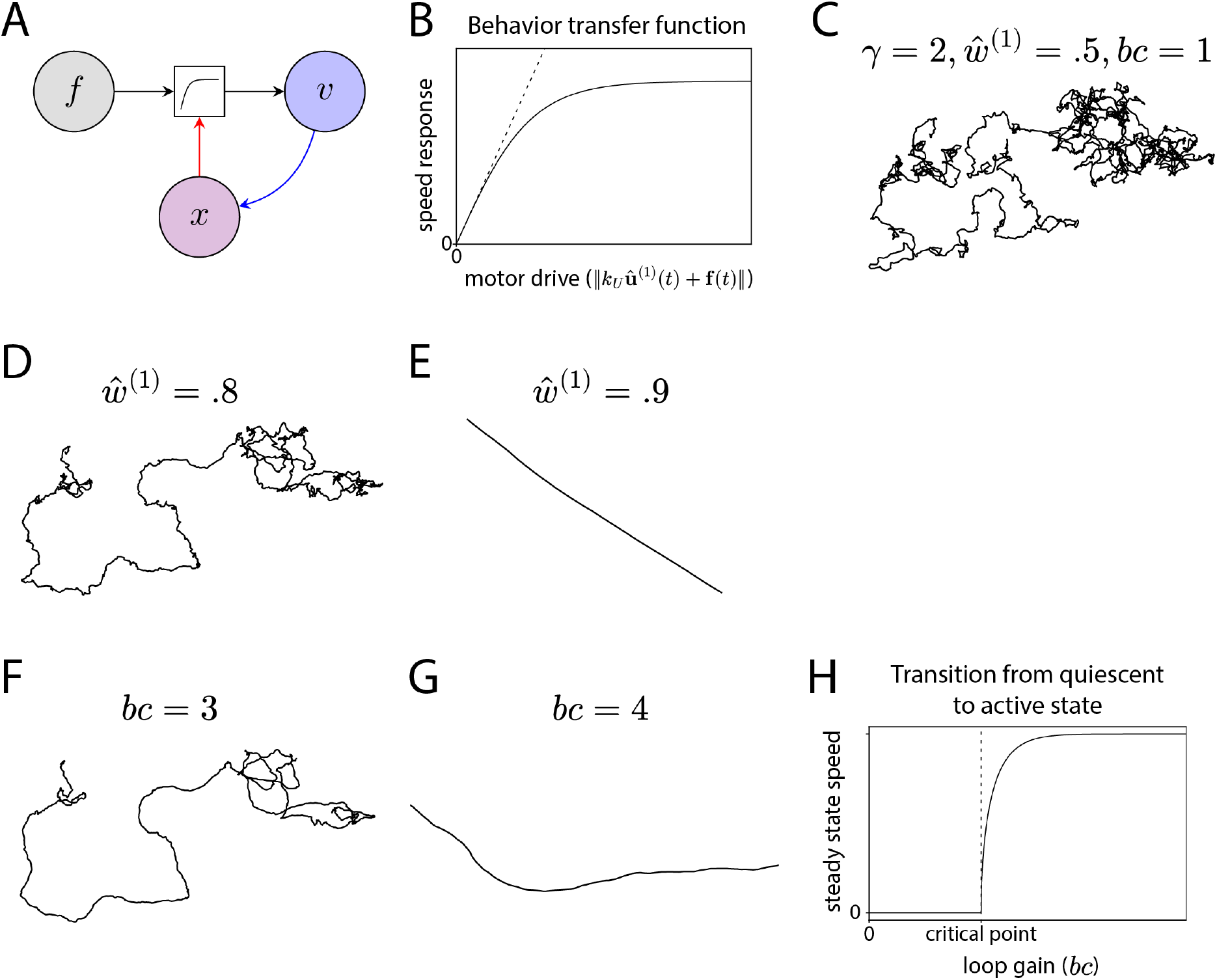
Agent behavior with a speed nonlinearity. (A) The motor drive that the agent inherits from the network and forcing function is now mediated by a nonlinear transfer function. (B) This transfer function is a saturating nonlinearity (hyperbolic tangent) that ensures that the agent’s speed remains bounded as the magnitude of the motor drive grows. (C) With this nonlinearity, as before, an agent with relatively weak coupling parameters produces a relatively meandering path. (D,E) In response to the same forcing, a network with stronger recurrence can generate a straighter path. (F,G) Increasing the gain of the control loop can also increase path straightness. (H) In the absence of forcing, the network undergoes a transition from a quiescent state lacking movement and neural activity to an active state in which activity is self-sustaining as the loop gain crosses a critical point.

The stability of this fixed point then depends on the leading eigenvalue *λ*_1_(**A**) of the coupling matrix, which is given by

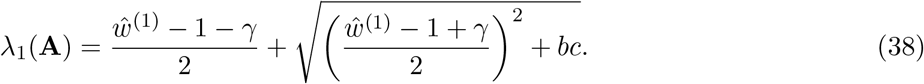

When the couplings are weak, this eigenvalue is negative and the fixed point is stable. However, as the eigenvalue crosses *λ*_1_(**A**) = 0, the quiescent state becomes unstable and self-sustaining activity emerges. Notably, this transition can be brought about by increasing the loop gain *bc*. Thus, even if neural activity and behavior are both dissipative in isolation, they can be sustained by strong sensorimotor feedback between the two. In other words, in this active state with strong feedback, neural activity is persistently maintained by the sensory input that continuously results from the circuit’s own motor output.

## Notes

### Competing Interest Statement

The authors have declared no competing interest.

